# *In vitro* evolution of caspofungin resistance in *Candidozyma auris* via *FKS1* hotspot I mutations results in moderate fitness trade-offs but no reduction in virulence

**DOI:** 10.1101/2024.12.18.629118

**Authors:** Zoe K. Ross, Sakinah Alsayegh, Yixin Zhao, Carol A. Munro, Alexander Lorenz

## Abstract

Resistance to echinocandins in the recently emerged pathogen *Candidozyma auris* (previously known as *Candida auris*) poses significant clinical challenges. This study reports the *in vitro* evolution of a caspofungin-sensitive *C. auris* clade II isolate to acquire caspofungin resistance. We sequenced the whole genome of the parental strain and its drug-resistant progeny. The resistant isolates harboured mutations in the *FKS1* gene altering a conserved residue (S639) also mutated in caspofungin-resistant clinical isolates; demonstrating that *FKS1-S639* mutations are sufficient to confer resistance to caspofungin.

Using these resistant strains, we examined cross-resistance & collateral sensitivity to other antifungal drugs and to cell wall stress, cell wall ultrastructure, biofilm formation, and virulence. As expected, both *FKS1-S639P* and *FKS1-S639Y* variants exhibited resistance not only to caspofungin but also to other echinocandins. Interestingly, these resistant strains showed moderate collateral sensitivity to certain azole antifungals and increased susceptibility to cell wall stress. While cell wall chitin content was slightly elevated in resistant isolates compared to sensitive ones, overall cell wall structure and phosphomannan levels remained unchanged. Biofilm formation capabilities were similar across all strains, and virulence in invertebrate models was unaffected by the *FKS1* mutations.

These findings indicate that while the evolution of caspofungin resistance through *FKS1* mutations does incur fitness trade-offs, it does not substantially compromise the pathogenic potential of *C. auris*. This underscores the complexities of evolution of antifungal resistance mechanisms in this pathogen and the impact on its biology.

## 1. Introduction

The human fungal pathogen *Candidozyma auris* – previously known as *Candida auris* (Liu et al., 2024) – has rapidly emerged as a global healthcare threat, causing nosocomial outbreaks with high mortality rates (Bravo Ruiz and Lorenz, 2021; Horton et al., 2023; Santana et al., 2024). Resistance to all classes of antifungals has been reported in the species and some isolates are pan-resistant, making effective treatment increasingly difficult (Lockhart et al., 2017; Ostrowsky et al., 2020). Around 5% of isolates are resistant to echinocandins, including caspofungin (CSP) (Chaabane et al., 2019; Lockhart et al., 2017). Resistance to echinocandins in a wide range of pathogenic yeast species is associated with mutations in *FKS1* and its paralog *FKS2*, which encode the catalytic subunit of β(1,3)-glucan synthase, or in its regulators (Daneshnia et al., 2023; Dudiuk et al., 2017; Hou et al., 2019; Khalifa et al., 2020; Martí- Carrizosa et al., 2015; Scott et al., 2023; Silva et al., 2023; Walker et al., 2010; Wiederhold, 2016). In *C. auris* isolates from clades I, III & IV, mutations in hotspot I of *FKS1* leading to substitution of serine at residue 639 by phenylalanine, tyrosine, or proline (S639F,-Y,-P) in the Fks1 protein have been correlated with naturally occurring echinocandin resistance (Biagi et al., 2019; Chow et al., 2020; Chowdhary et al., 2018; Kordalewska et al., 2018). Indeed, *C. auris* strains isolated during hospital outbreaks in London and Shenyang that were found to be echinocandin-resistant had *FKS1* hotspot mutations (Rhodes et al., 2018; Tian et al., 2024). Importantly, *in vitro* and *in vivo* evolution experiments for echinocandin resistance also resulted in S639 mutations (Carolus et al., 2021; Hirayama et al., 2023). Furthermore, variants and engineered mutations outside the hotspots of *FKS1* have also been reported to reduce susceptibility to echinocandins in *Candida parapsilosis* and *C. auris* (Carolus et al., 2021; Daneshnia et al., 2023; Hirayama et al., 2023; Kiyohara et al., 2023; Kordalewska et al., 2023). However, *C. auris* clade II strains rarely cause systemic candidiasis and are predominantly reported from ear infections, tend to be sensitive to antifungal drugs except fluconazole, and, so far, did not evolve S639 *FKS1* hotspot I mutations (Carolus et al., 2021; Chow et al., 2020; Hirayama et al., 2023; Kwon et al., 2019; Welsh et al., 2019).

Here, we *in vitro* evolved CSP-resistant (CSP^R^) strains from the CSP-sensitive (CSP^S^) *C. auris* clade II isolate B11220. The CSP^R^ derivatives and the CSP^S^ control strain had their genomes sequenced to get an unbiased view of the genomic changes leading to CSP resistance. For the first time we could show that clade II isolates can evolve resistance to echinocandins via *FKS1* hotspot I mutations. This has not been achieved previously in *in vitro* and *in vivo* approaches (Carolus et al., 2021; Hirayama et al., 2023). Our study also emphasises that despite the unique genomic structure and features of clade II *C. auris* strains (Bravo Ruiz et al., 2019; Muñoz et al., 2019, 2018), they respond to antifungal drug treatment in a similar way compared to strains from other clades. We observed subtle changes to cell wall biology and moderate fitness trade-offs in CSP^R^ *FKS1* variant strains compared to CSP^S^ control strains. Overall, this demonstrates that evolution of CSP resistance via *FKS1* hotspot I mutation despite having some fitness trade-offs does not obviously impact the pathogenicity potential of *C. auris*.

## 2. Materials & Methods

### 2.1. *In vitro* evolution

*Candidozyma auris* strain B11220 (Table S1) was incubated in RPMI-1640 (RPMI) [10.4 g/l RPMI-1640 powder without phenol red (Merck KGaA, Darmstadt, Germany), 1.8% D-Glucose (ThermoFisher Scientific, Waltham, MA, USA), 0.165 M 3-(N-morpholino)propanesulfonic acid (Melford, Ipswich, UK) adjusted to pH 7 by 1 M NaOH (Sigma-Aldrich, St. Louis, MO, USA)] liquid medium with or without 0.125 μg/ml caspofungin (CSP) (Sigma-Aldrich) for 24 h at 30°C. These cultures were passaged 1:10 into fresh liquid medium containing the same concentration of CSP and incubated as above. Cells were harvested at 3,500 ×*g* for 5 min and washed in sterile ddH_2_O before being spread onto RPMI agar plates containing 0, 0.125 or 0.5 μg/ml CSP. The plates were incubated for 4 days at 30°C. Colonies were picked from 0 and 0.5 μg/ml CSP plates and resuspended in RPMI broth with or without 0.5 μg/ml CSP. These cultures were passaged once more into fresh RPMI broth containing 0, 0.5, or 1 μg/ml CSP, and incubated for a further 24 h. Finally, cells were harvested, washed, and spread onto Yeast extract Peptone Dextrose (YPD) [1% yeast extract, 2% mycological peptone, 2% glucose, 2% agar (all Oxoid Ltd., Basingstoke, UK)] agar plates. Overnight cultures in YPD broth from the YPD agar plates were then tested for CSP sensitivity in comparison to the parental strain. Two isolates were tested from each of two *in vitro* evolution procedures – UACa141 and UACa142 from one, and UACa143 and UACa144 from the other (see Tables 1, S1).

### 2.2. Whole-genome sequencing and bioinformatics

Sodium azide (final concentration 0.1%, Sigma-Aldrich) was added to overnight *C. auris* cultures before centrifugation (4 min, 1,050 ×*g*). After removing the supernatant, the cell pellet was frozen at-20°C for 18-24 h. The pellet was resuspended in spheroplasting buffer [1 M sorbitol, 50 mM KPO_4_ pH 7, 10 mM EDTA pH 7.5, 1/100^th^ volume β-mercaptoethanol (Sigma-Aldrich), 20T Zymolyase (2.5 mg/ml, AMS Biotechnology, Abingdon, UK)] and incubated at 37°C for 15 min. Spheroplasts were harvested by centrifugation (4 min at 2,000 ×*g*) and resuspended in Guanidine solution [4.5 M Guanidine-HCl (Sigma-Aldrich), 0.1 M EDTA, 0.15 M NaCl, 0.05% N-lauroyl sarcosine sodium salt (Sigma-Aldrich)]. The spheroplasts were then incubated at 65°C for 20 min, cooled on ice, and mixed with 1 volume of 100% ethanol. After further incubation on ice (20 min), samples were centrifuged (15 min, 3,500 ×*g*). The pellets were incubated in RNAse solution (10× TE pH 8, 50 µg/ml RNase A) for 1 hr at 37°C before proteinase-K solution (20 mg/ml in 20mM CaCl_2_, 10 mM Tris-HCl pH 7.5, 50% glycerol) was added for 1 hr incubation at 50°C. DNA was then extracted with 1 volume of phenol:chloroform:isoamyl alcohol (25:24:1, Sigma-Aldrich). After mixing by inversion, the samples were centrifuged (10 min, 12,000 ×*g*). The top aqueous layer was transferred to a fresh reaction tube and the phenol/chloroform extraction was repeated. To precipitate the DNA, 1/20^th^ volume of 3 M sodium acetate pH 5.2 and 2 volumes 100% ethanol were added. After mixing, the samples were centrifuged (5 min, 12,000 ×*g*), and the DNA pellets were rinsed in 70% ethanol. DNA pellets were air-dried and resuspended in ddH_2_O. The concentration and quality of the DNA samples were determined on a NanoDrop™ 2000 spectrophotometer (ThermoFisher Scientific). Whole-genome sequencing was performed on an Illumina NextSeq500 with a Nextseq mid-output 300c v2.5 flowcell (Illumina Inc., San Diego, CA, USA) at the University of Aberdeen Centre for Genome Enabled Biology and Medicine (CGEBM). Illumina-sequencing reads were aligned to the B11220 genome (GCA_003013715.2_ASM301371v2_genomic.fna) (Muñoz et al., 2018) using BWA-MEM v0.7.12 (Li and Durbin, 2010) and processed with Samtools v0.1.19 view, sort, rmdup and index (Li et al., 2009). SNPs were detected using Pilon v1.22 (Walker et al., 2014) and the resulting variant call format (VCF) file was filtered for genotype 1/1 only. Low coverage (less than 10% of mean coverage), ambiguous positions and deletions were removed. The reference genome (B11220, GCA_003013715.2_ASM301371v2_genomic.fna) (Muñoz et al., 2018) was annotated with Augustus v3.3.1 (Stanke et al., 2004) *ab initio* gene prediction software and VCF annotator (http://vcfannotator.sourceforge.net) was used to predict the effect of the SNPs called on the annotated genes.

### 2.3. Caspofungin Epsilometer tests (Etests) and spot assays

Overnight *C. auris* cultures in YPD broth were harvested by centrifugation, washed twice and resuspended in sterile ddH_2_O. Cell density was adjusted to ∼1 × 10^7^ cells/ml, and 1 × 10^6^ cells were spread onto RPMI agar plates (bioMérieux, Marcy-l’Étoile, France). A CSP Etest strip (Liofilchem S.r.l., Roseto degli Abruzzi, Italy) was then placed with sterile forceps onto the agar plates, avoiding air bubbles underneath the strip. Plates were incubated at 37°C and 5% CO_2_ in a humid incubator for 24 h. CSP MICs were read visually from the strip.

For spot assays, overnight *C. auris* cultures in RPMI broth were harvested by centrifugation, washed twice and resuspended in sterile ddH_2_O, cell density was adjusted to 1 × 10^7^ cells/ml. Spot assays were performed on solid RPMI medium as previously described (Bravo Ruiz et al., 2020). Antifungal drugs and compounds inducing cell wall & osmotic stress were sourced commercially. Antifungal drugs – caspofungin (CSP), anidulafungin (AFG), micafungin (MFG), 5-fluorocytosine (5-FC), itraconazole (ICZ), posaconazole (PSZ), voriconazole (VCZ) (all Sigma-Aldrich), and fluconazole (FCZ; Discovery Fine Chemicals Ltd, Ferndown, UK) – were dissolved in DMSO (Sigma-Aldrich) to give appropriate stock solutions. Cell wall & osmotic stress compounds – Calcofluor White (CFW; Sigma-Aldrich), Congo Red (CR; Sigma-Aldrich), sodium chloride (NaCl; ThermoFisher Scientific), sorbitol (Sigma-Aldrich) – were either dissolved in sterile ddH_2_O to produce stock solutions or directly added to the medium before sterilisation.

### 2.4. Flow cytometry to measure chitin content of cell walls

Strains were inoculated in RPMI medium and incubated shaking (200 rpm) at 30°C for 16 hours. Overnight cultures were washed three times with 1× PBS and fixed in 4% paraformaldehyde (ThermoFisher Scientific) at 37°C for 15 minutes. Fixed cells were washed three time with 1× PBS and stained in the dark with 10 μg/ml Calcofluor White (CFW) at room temperature for 1 h. Cells were washed three times with 1× PBS again before detection on a flow cytometer. An accelerating voltage of 200 kV for VL1 (excitation laser 405 nm and emission filter 440/50) was used to detect CFW signals on an Attune flow cytometer. The samples were gated for singlets using forward scatter height and forward scatter area; around 50,000 single cells were measured for each sample.

### 2.5. Cell wall thickness measurements and phosphomannan content quantification

Cells were grown for 16 hours in RPMI medium and harvested by centrifugation (2,500 ×*g*, 5 minutes). Excess cell culture medium was removed before fixation and preparation. High-pressure freezing on a Leica Empact 2/RTS high-pressure freezer was used to fix samples by the Microscopy and Histology Core Facility at the University of Aberdeen following a published protocol (Milne and Walker, 2022). Samples were imaged using a JEOL 1400 plus transmission electron microscope with an AMT UltraVUE camera. Fiji (ImageJ version 1.53f51) (Schindelin et al., 2012) was used to process images and measure cell wall thickness.

Phosphomannan content was determined by Alcian blue staining (Hobson et al., 2004). *C. auris* isolates and the controls – *Candida albicans* wild type (SC5314) and the *C. albicans mnn4*Δ/*mnn4*Δ mutant (see Table S1) – were incubated shaking (200 rpm) for 24 hr in 10 ml RPMI broth at 30°C. The cells were washed twice with sterile ddH_2_O via centrifugation (2,500 ×*g*, 5 minutes) and finally resuspended in 10 ml sterile ddH_2_O. The cell density was determined and adjusted to 10^8^ cells/ml in water and samples were transferred to 1.5-ml reaction tubes. Cells were centrifuged at 5,000 ×*g* for 1 min, the supernatant was discarded, and the cell pellet was re-suspended in 1 ml of Alcian Blue (Alcian Blue 8GX, Merck KGaA, Darmstadt, Germany) solution (30 µg/ml in 0.02M HCl). After incubation at room temperature for 10 min, the cells were pelleted at 5,000 ×*g* for 1 min. 200 µl of the supernatant from each sample was transferred to a well of a 96-well plate. The absorbance at 620 nm was measured spectrometrically on a VERSAmax tunable microplate reader (Molecular Devices LLC, San Jose, CA, USA). Concentration of Alcian Blue was determined by referring to a standard curve. The amount of Alcian Blue bound to the cell wall in pg/cell was calculated according to Hobson and co-workers (Hobson et al., 2004).

### 2.6. Biofilm mass and viability testing

Biofilms were established in flat-bottom 96-well plates (ThermoFisher Scientific) from overnight *C. auris* cultures in RPMI. 200 µl of *C. auris* cells at an OD_600nm_ of 0.5 in fresh RPMI were added to each well, and incubated at 30°C. The growth medium was removed, and non-adherent cells were gently washed away using 200 µl of 1× PBS by pipetting along the sides of the wells. The 96-well plate was then left to dry for 24 hours on the benchtop. Crystal Violet assay was performed following an established protocol (Mowat et al., 2007; Peeters et al., 2008) to quantify biofilm mass. To determine cell viability within biofilm the resazurin-based AlamarBlue assay (ThermoFisher Scientific) was used according to instruction of the manufacturer (see also Brown et al., 2022). All absorbance measurements were performed on a VERSAmax tunable microplate reader (Molecular Devices LLC).

### 2.7. Galleria mellonella survival assay

Virulence testing in the invertebrate infection model *Galleria mellonella* was in essence performed as previously described (Pelletier et al., 2024). *G. mellonella* larvae (UK Waxworms Ltd., Sheffield, UK) were inoculated within two days of receipt. Larvae within a weight range of 280 – 350 mg without dark splotches were organized into groups of 10 per petri dish and incubated in the dark at room temperature for 24 hrs.

*C. auris* strains were picked as single colonies from a Sabouraud dextrose (SabDex) plate [1% mycological peptone (Oxoid Ltd.), 4% D-glucose (ThermoFisher Scientific), 2% agar (Formedium Ltd., Swaffham, UK)] plate and grown for 16 hours in RPMI broth. Medium was removed by centrifugation (2,500 ×*g*, 5 minutes), cells were washed twice with ddH_2_O by centrifugation (2,500 ×*g*, 5 minutes) between each resuspension. Finally, cells were suspended in 1× PBS to give a final inoculum of 10 µl containing 1 × 10^6^ cells. The inoculum was given via the last left proleg with a 0.5 ml 29G Micro-Fine U-100 insulin injection unit (BD Medical, Franklin Lakes, NJ, USA).

Larvae were incubated for 6 days at 37°C. Every 24 hours larvae were examined and deemed dead when they no longer responded to physical stimuli. Each experiment had a non-injected control group, and a control group injected with 1× PBS only; survival in the control groups over the 6-day incubation period was 100%. Results were pooled across two independent experiments and visualised with a Kaplan-Meier survival plot.

### 2.8. Data visualization and statistics

Kaplan-Meier survival plots (Pelletier et al., 2024) and box plots (Lorenz et al., 2014) were generated with R (version Rx64, 4.4.0) (https://www.r-project.org/). Bar charts showing individual data points were made in Excel (version 2407, Microsoft Corp., Redmond, WA, USA). Images were processed and assembled in Adobe Photoshop (version 25.11.0) and Adobe Illustrator (version 28.7) (Adobe Inc., San Jose, CA, USA).

Statistical analysis (Kruskal-Wallis test, Dunn’s test with Bonferroni correction, ANOVA, Tukey’s HSD, and log-rank test) was performed in R (version Rx64, 4.4.0).

## 3. Results & Discussion

### 3.1. *In vitro* evolution of *C. auris* clade II strain B11220 to caspofungin resistance

*C. auris* clade II strains tend to be susceptible to most clinically used antifungal drugs and apparently rarely cause systemic candidiasis (Chow et al., 2020; Kwon et al., 2019; Welsh et al., 2019). We performed an *in vitro* evolution experiment with the CSP^S^ clade II strain B11220 to understand whether clade II isolates are deficient in developing antifungal resistance. B11220 was incubated in RPMI broth with 0.125 μg/ml CSP for 2 days. Cells were then spread onto solid RPMI containing 0.125 or 0.5 μg/ml CSP and two colonies each were picked at random from two independent *in vitro* evolution assays for further passaging (2 days) in an increased concentration of CSP (0.5 or 1 μg/ml CSP). In parallel, B11220 was passaged in RPMI without CSP, a single isolate (UACa140) was selected as a control strain. UACa140 remained sensitive to CSP in an Etest (Figs. 1, S1), whereas the four isolates that had been passaged in medium containing CSP were now CSP^R^ (UACa141, UACa142, UACa143, UACa144) (Figs. 1, S1). After two weeks growth on RPMI lacking CSP, the CSP^R^ isolates (UACa141-UACa144) had maintained resistance to CSP (Fig. S2), indicating that these *in vitro* evolved strains are resistant likely due to genetic mutation rather than just having adapted to higher CSP concentrations (e.g. by cell wall remodelling).

**Figure 1.**
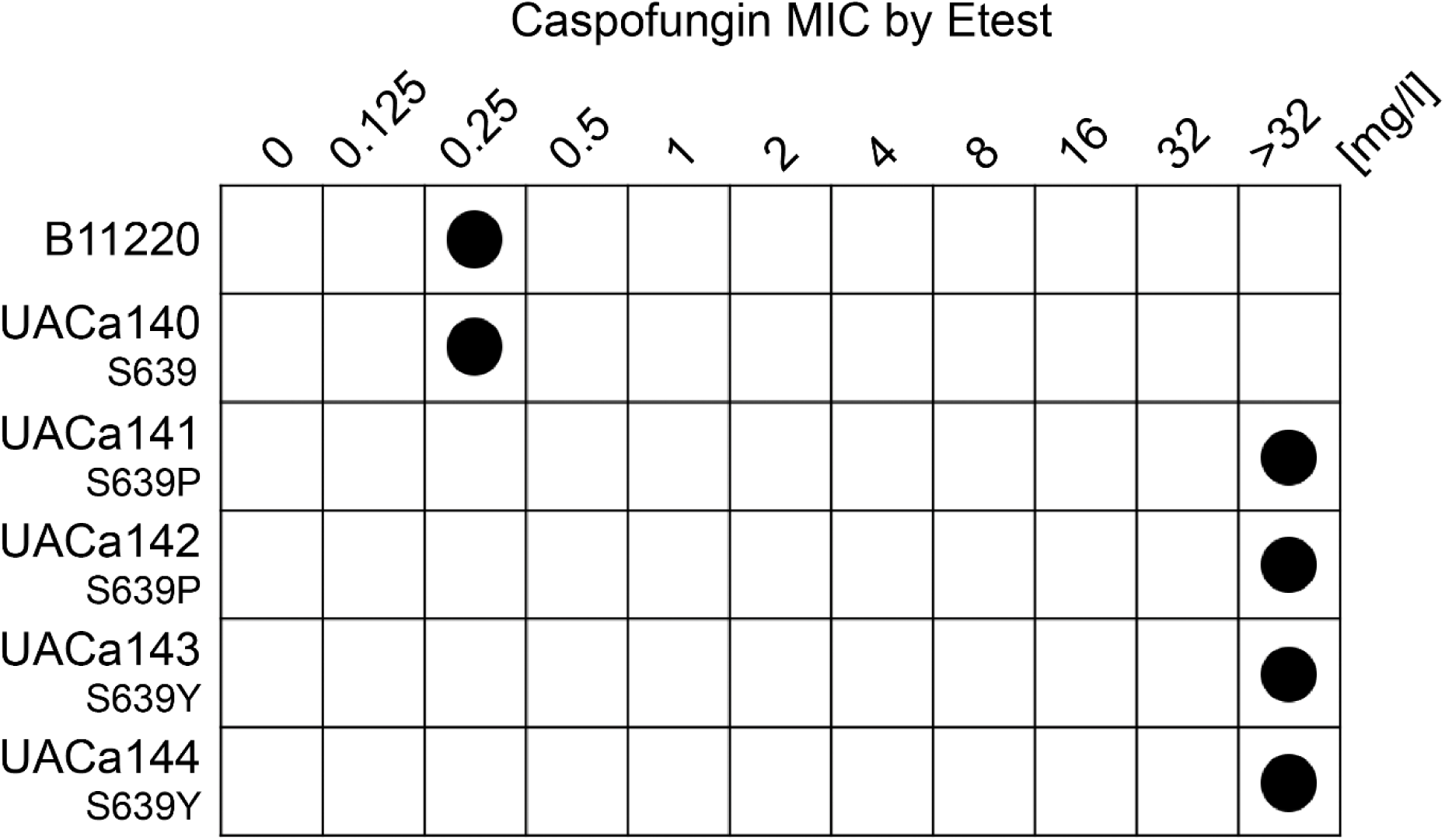
Caspofungin MICs of parental strain (B11220) and *in vitro* evolved isolates as determined by Etest (see also Figs. S1, S2). *FKS1* variants are indicated underneath strain number (see Table 1 for details).

**Table 1.**
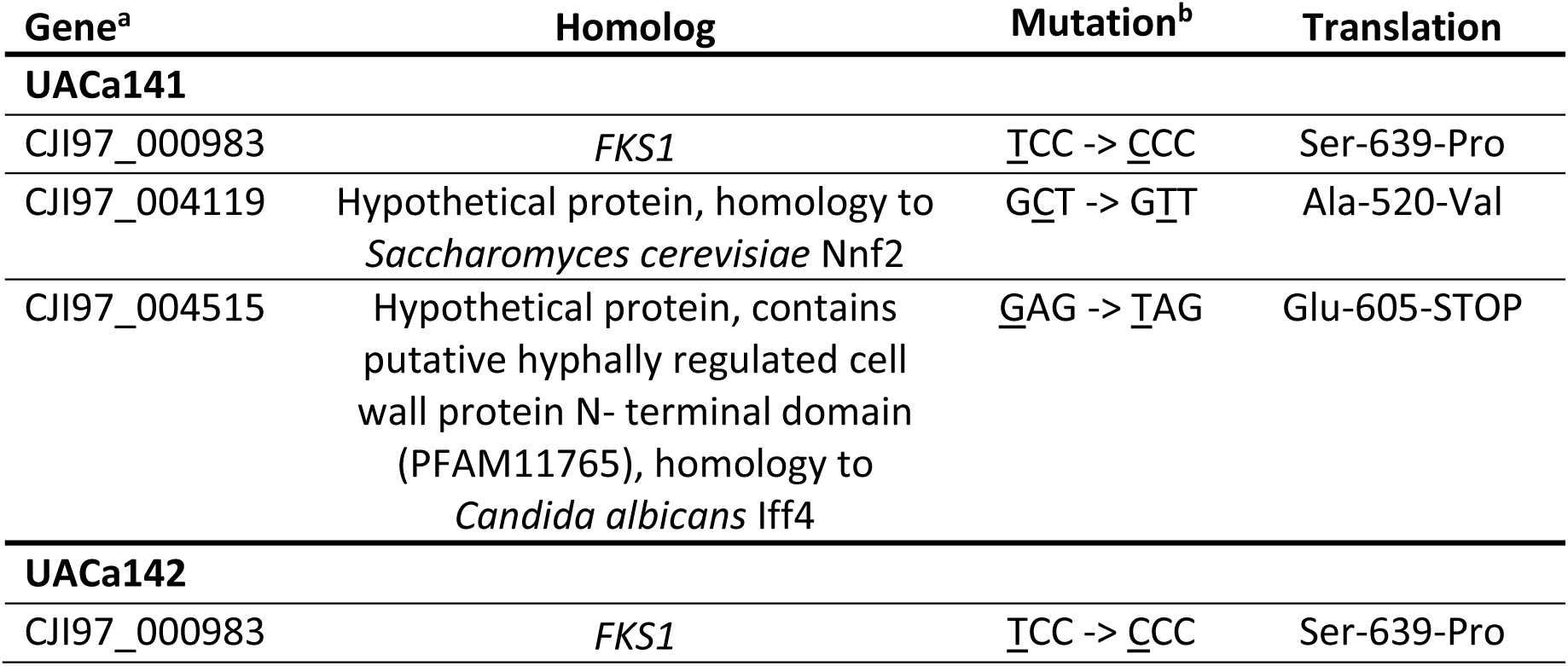
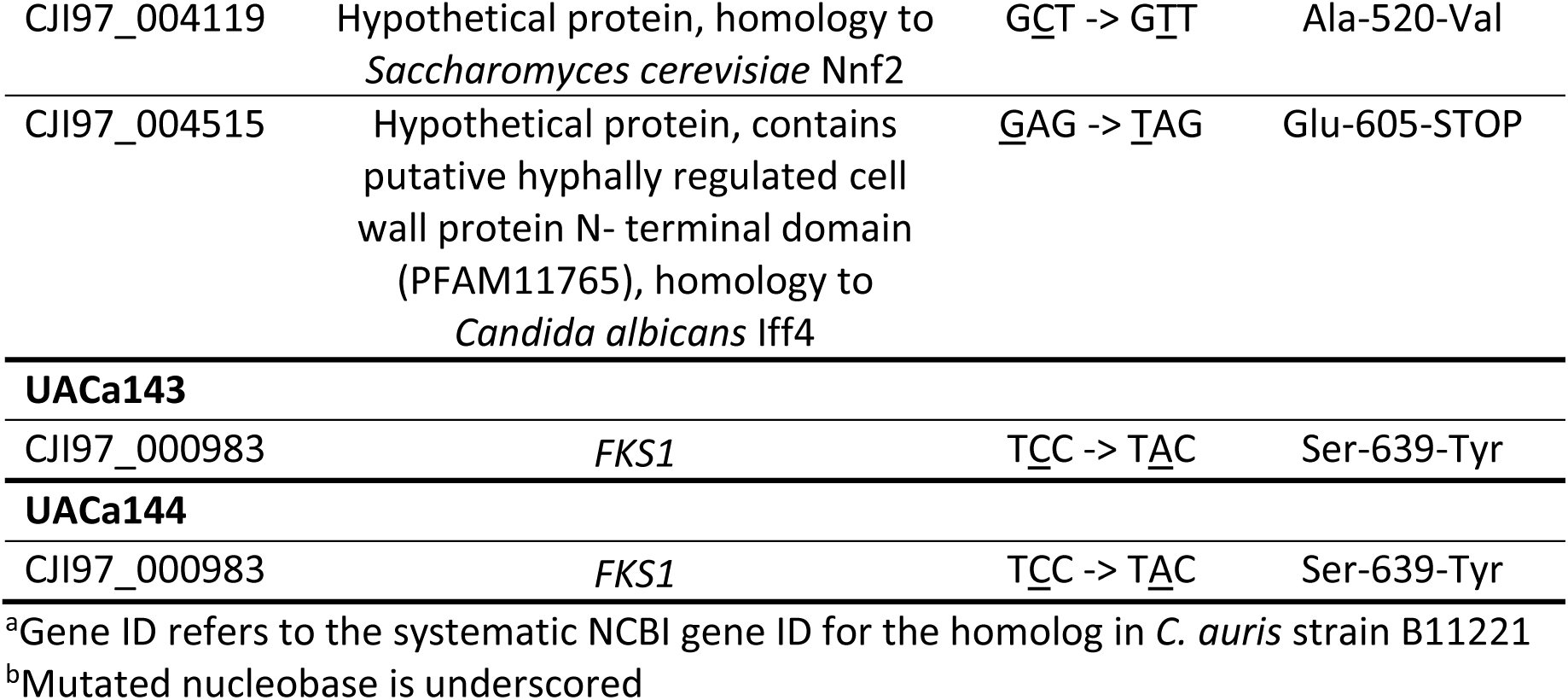
SNPs identified in coding regions of caspofungin-resistant isolates.

### 3.2. Whole-genome sequencing of *in vitro* evolved strains

To gather an unbiased view of all genomic DNA sequence changes during the *in vitro* evolution experiment of B11220, the CSP^R^ isolates (UACa141-UACa144) and the sensitive control strain UACa140 were genome-sequenced on an Illumina NextSeq platform (ENA Accession No. <u>PRJEB40741</u>). The Illumina sequencing reads were aligned to the published B11220 genome (GCA_003013715.2_ASM301371v2_genomic.fna) (Muñoz et al., 2018) and variants were called. The isolate passaged without CSP (UACa140) had 30 SNPs compared to B11220, which were all located in intergenic regions. Between B11220 and the CSP^R^ isolates (UACa141-UACa144), the number of SNPs ranged from 35 to 39, with one to four SNPs within genes and the remaining SNPS (>30) in non-coding regions. The B11220 genome was annotated, and this annotation was used to predict function of the genes containing SNPs. These SNPs in the CSP^R^ isolates involved only four different genes. One SNP found in two isolates UACa141 and UACa142 was located in the intron region of a putative histone acetyltransferase homolog, without affecting splice sites. These isolates also had SNPs in the coding regions of a putative homolog of *Saccharomyces cerevisiae NNF2*, and of a putative homolog of *C. albicans IFF4* (Table 1). Notably, the *FKS1* gene was mutated in all four CSP^R^ isolates (Table 1). UACa141 and UACa142 arose from the same *in vitro* evolution experiment, and UACa143 and UACa144, arose from a separate evolution lineage, both pairs had the same SNPs within genes. Therefore, we cannot exclude the possibility that these are pairs of clones. Having said that, there are differences in intergenic SNPs between UACa141 and UACa142, as well as UACa143 and UACa144. The mutation affects S639 of Fks1 in all cases (Table 1), which is the same hotspot residue that is mutated in CSP^R^ clinical isolates from other clades (Rhodes et al., 2018). The mutations cause the serine residue at position 639 to be translated to tyrosine or proline, both of which are hydrophobic. Our results suggest that a mutation leading to a hydrophobic residue at position 639 of Fks1 is sufficient to provide resistance to CSP in *C. auris*. This is consistent with previous findings of CSP resistance caused by S639F, S639Y or S639P residue change in *C. auris* clade I, III & IV isolates (Chow et al., 2020; Chowdhary et al., 2018; Tian et al., 2024), by S638P and S638Y residue changes in clinical *Clavispora lusitaniae* isolates (Asner et al., 2015), and S645P and S645Y residue changes in *C. albicans* (Desnos-Ollivier et al., 2008). Overall, this is the first time that a mutation of this hotspot I site resulting in a tyrosine or proline amino acid change has been described for a *C. auris* clade II isolate (Table 1). Another *in vitro* evolution experiment of the clade II strain B11220 resulted in mutations other than the *FKS1-S639* variants reported here (Carolus et al., 2021). Interestingly, attempts to *in vivo* evolve a clade II strain (JCM15448) to echinocandin resistance in a murine gut colonisation model failed altogether (Hirayama et al., 2023). Nevertheless, this report and previous studies (Carolus et al., 2021; Hirayama et al., 2023) verify a direct link between CSP selection pressure, echinocandin resistance, and *FKS1* mutations in *C. auris*. Our data corroborates that clade II isolates, which tend to be susceptible to most antifungals (Chow et al., 2020; Kwon et al., 2019), can rapidly gain resistance when antifungal selection pressure is applied (Carolus et al., 2024a, 2021).

### 3.3. *FKS1-S639* variant strains are resistant to echinocandins but show collateral sensitivity to some azole drugs and are sensitive to cell wall stress

To test how *C. auris FKS1-S639P/Y* variants impact on broader antimicrobial resistance, we compared the growth of CSP^S^ *FKS1-S639* isolates with that of CSP^R^ *FKS1-S639P/Y* strains on solid RPMI medium containing a series of antifungal drugs. As expected, *FKS1-S639P* and *FKS1-S639Y* strains were not only resistant to CSP but also considerably less susceptible to other echinocandin antifungals, represented by anidulafungin (AFG) and micafungin (MFG) (Fig. 2A). This is consistent with all echinocandin drugs inhibiting the β-(1,3)-glucan synthase Fks1 (Perlin, 2015).

**Figure 2.**
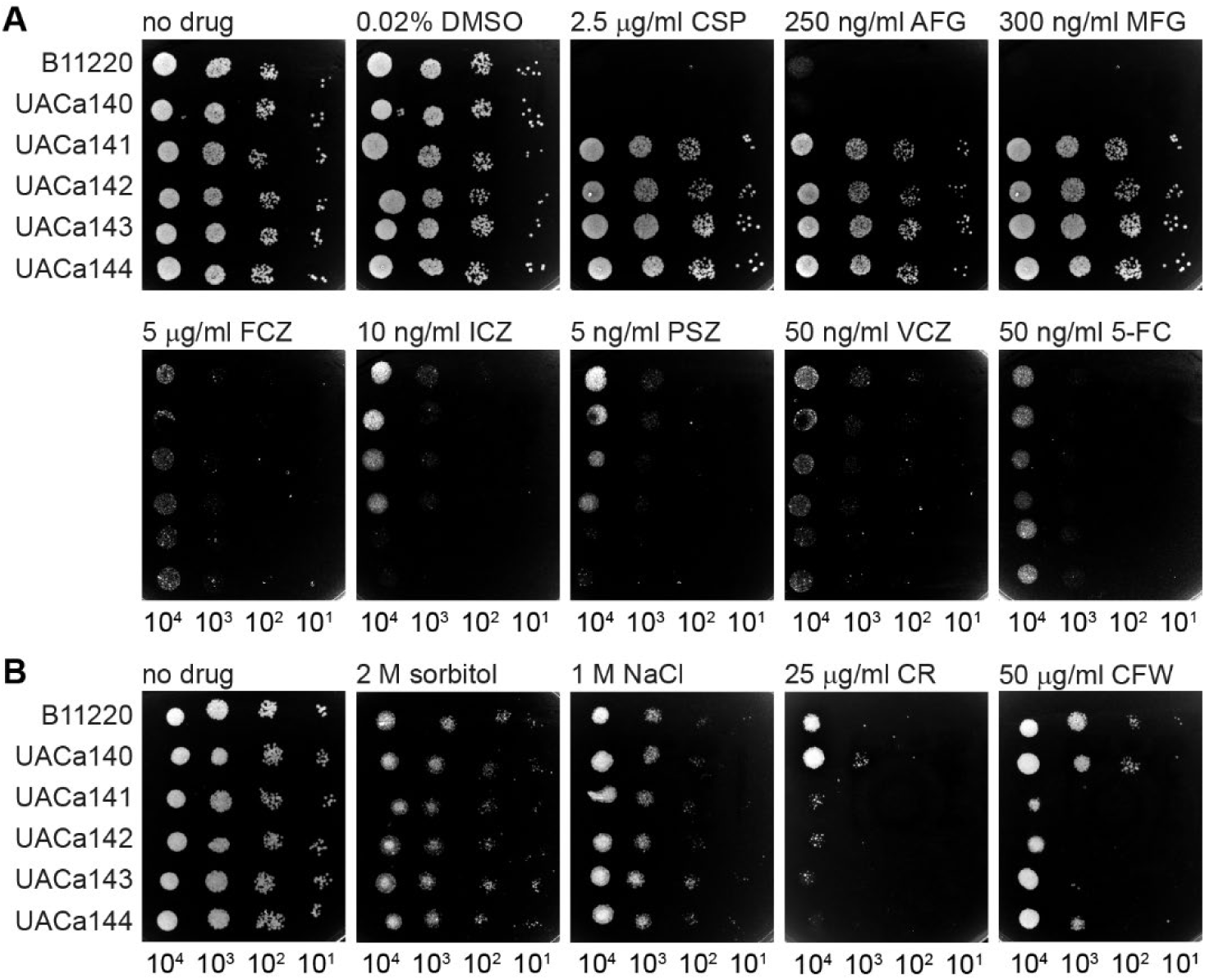
The response of *FKS1-S639* variant *Candidozyma auris* clade II strains to (**A**) antifungal drugs and (**B**) cell wall & osmotic stress. Growth analysis by spot assays of CSP^S^ strains B11220 & UACa140, and CSPR strains UACa141, UACa142, UACa143 & UACa144 in the presence and absence of treatments. 10-fold serial dilution of *C. auris* cells were grown on RPMI plates containing the indicated compound for 72 hours at 30°C. Dimethylsulfoxide (DMSO), caspofungin (CSP), anidulafungin (AFG), micafungin (MFG), fluconazole (FCZ), itraconazole (ICZ), posaconazole (PSZ), voriconazole (VCZ), 5-fluorocytosine (5-FC), sodium chloride (NaCl), Congo Red (CR), Calcofluor White (CFW).

Intriguingly, the CSP^R^ *FKS1-S639P/Y* variant strains showed some collateral sensitivity to some azoles, especially itraconazole (ICZ) and posaconazole (PSZ), compared to the CSP^S^ control strains (Fig. 2A); this is particularly obvious in the *FKS1-S639Y* variant strains UACa143 and UACa144. The relative lipophilicity of ICZ and PSZ in comparison to fluconazole (FCZ) could play a role here. However, this would not explain the lack of a differential response between CSP^S^ and CSP^R^ strains to voriconazole (VCZ) (Fig. 2A) which is also lipophilic (Bellmann and Smuszkiewicz, 2017). The diverging susceptibility to ICZ and PSZ of *FKS1-S639P* and of *FKS1-S639Y* isolates points to potential differences in the cell wall and/or membrane of these variants. But, this will likely be an indirect effect and we also cannot exclude that this might be influenced by other genetic differences between these strain pairs (see Table 1).

The susceptibility to 5-fluorocytosin (flucytosine, 5-FC) remains unaltered in *FKS1-S639P/Y* variant strains (Fig. 2A). In contrast to our observation, Carolus and colleagues reported collateral sensitivity to 5-FC in *in vitro* evolved CSP^R^ derivatives of a *C. auris* clade II strain (Carolus et al., 2024a). As the *FKS1* variant status of the CSP^R^ strains in Carolus et al. (2024a) has not been determined, a likely explanation for this discrepancy is that the genetic changes in their CSP^R^ strains are not affecting *FKS1*, or at least do not comprise changes in hotspot I of *FKS1*.

To understand how *FKS1-S639P/Y* variants might influence the integrity of the cell wall, we tested whether *FKS1* hotspot I status influenced susceptibility to cell wall stress (Ram and Klis, 2006) and high levels of hyperosmotic treatment (Hohmann and Mager, 2003).

No difference in growth was observed between CSP^S^ *FKS1-S639* strains and CSP^R^ *FKS1- S639P/Y* variants when exposed to high concentrations of sorbitol (hyperosmotic stress) or sodium chloride (NaCl, hyperosmotic/cationic stress) (Fig. 2B).

*FKS1-S639P/Y* variant isolates were considerably more sensitive to cell wall-specific stress induced by Congo Red (CR) and Calcofluor White (CFW) (Fig. 2B). Intriguingly, the response to CFW shows a difference between *FKS1-S639P* and *FKS1-S639Y* strains again highlighting the possibility of different consequences of these point mutations for the functionality of the resulting protein (see above, regarding the response to some azole drugs).

Overall, the spot assays demonstrate genuine differences in cellular behaviour of the CSP^S^ *FKS1-S639* strains and CSP^R^ *FKS1-S639P/Y* variants, indicating that point mutations that change residue S639 of Fks1 lead to resistance against echinocandins but sensitizes the cells to some azole drugs and to cell wall stress (Fig. 2). This implies that there is a fitness trade-off in *C. auris* which has evolved resistance to CSP via mutation of *FKS1*, similar to what has been observed in *C. albicans FKS1-S645* variants (Ben-Ami et al., 2011).

### 3.4. *FKS1* variants have a higher cell wall chitin content but do not show altered cell wall ultrastructure and phosphomannan levels

Yeast pathogens of humans – including *Candida* spp., *Meyerozyma guilliermondii*, *Candidozyma auris* – can adapt to echinocandin treatment by increasing cell wall chitin (Ben-Ami et al., 2011; Lara-Aguilar et al., 2021; Lee et al., 2012; Walker et al., 2013). This likely is the result of a compensatory mechanism to strengthen the cell wall when the β-(1,3)-glucan synthase Fks1 is inhibited by echinocandin treatment (Walker et al., 2010, 2008). *FKS1* variant strains of several pathogenic yeast species seem to have a higher baseline level of cell wall chitin and do not gain more chitin when treated with CSP (Ben-Ami et al., 2011; Walker et al., 2013). The increase in chitin level after CSP treatment is more pronounced in *C. albicans* than in *C. auris* (Lara-Aguilar et al., 2021). Therefore, we explored whether the *C. auris FKS1-S639P/Y* variant strains show differences in cell wall ultrastructure and biochemistry compared to the CSP^S^ *FKS1-S639* strains.

We measured the total cell wall chitin content by flow cytometry using CFW as a chitin-specific stain on fixed *C. auris* cells. *FKS1-S639Y* variant strains, UACa143 & UACa144, had significantly higher chitin content compared to the CSP^S^ strains B11220 and UACa140 (Figs. 3A, S3; Tables S2, S3). *FKS1-S639P* variant strains, UACa141 & UACa142, had moderately higher chitin content compared to the CSP^S^ strain B11220, statistical significance was only achieved in one of the flow cytometry runs for UACa141 and UACa142 each (Figs. 3A, S3; Tables S2, S3). The *FKS1-S639P* variant strains did not have a significantly higher cell wall chitin content in comparison to the CSP^S^ strain UACa140 (Figs. 3A, S3; Tables S2, S3). However, the two CSP^S^ strains B11220 and UACa140 did not differ significantly from each other in both runs (Figs. 3A, S3; Tables S2, S3).

**Figure 3.**
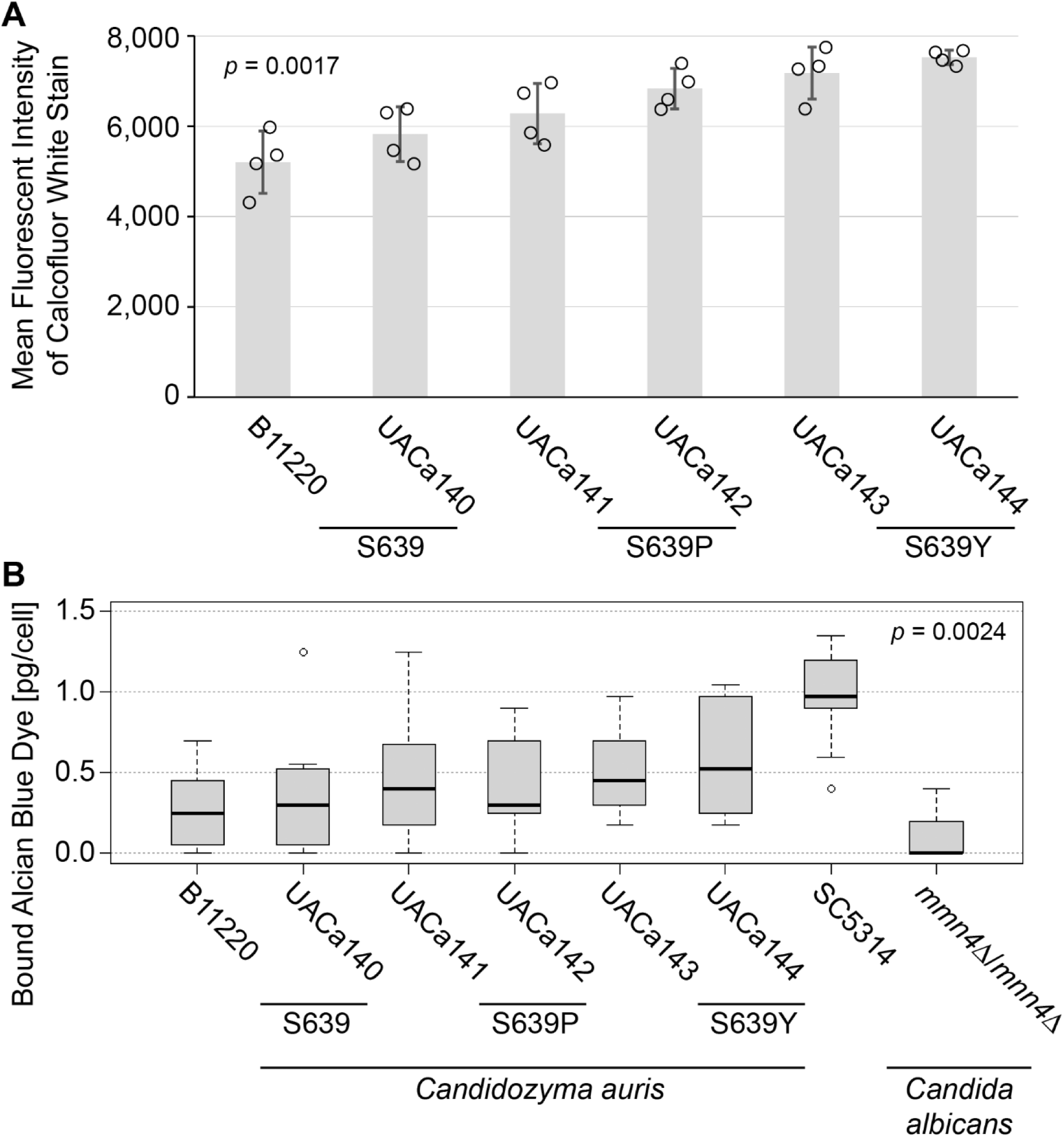
Cells of CSP^R^ strains have a higher chitin but largely unchanged phosphomannan content. (**A**) Bar chart showing mean fluorescent intensity of Calcofluor White staining of the indicated *C. auris* strains determined by flow cytometry (n = 4, ∼50,000 cells measured in each sample) *p*-value from an ANOVA, for *post-hoc* analysis with Tukey’s HSD see Table S2. An independent run is shown in Fig. S3, for representative examples of the gating strategy see Fig. S4. (**B**) Boxplot showing Alcian Blue staining [pg/cell] of the indicated *C. auris* and *C. albicans* strains (n = 9 each). *p*-value from a Kruskal-Wallis test, for *post-hoc* analysis with a Dunn’s test using a Bonferroni correction see Table S4.

Negatively charged glycosylated mannoproteins form the outermost shell of the cell wall of several fungi including *C. auris* (Alvarado et al., 2024; Bruno et al., 2020; Gow and Lenardon, 2023). Phosphate groups linking the mannose chains provide the negative charge (Gow and Lenardon, 2023) and influence how efficiently a given fungus is recognised by the immune system and also impacts the binding of antimicrobial peptides (Harris et al., 2009; McKenzie et al., 2010). Interestingly, *C. albicans* mutants adapted to caspofungin but not harbouring *FKS1* variants displayed a lowered phosphomannan content (Sah et al., 2022). In this experiment we also included two *C. albicans* strains as controls; the laboratory strain SC5314 as a positive one and the *mnn4*Δ/*mnn4*Δ deletion mutant in the same strain background as a negative one (Hobson et al., 2004). Phosphomannan content of CSP^S^ and CSP^R^ *C. auris* strains was not significantly different from each other, although the CSP^R^ strains seem to have slightly higher phosphomannan content per cell (Fig. 3C, Table S4). Generally, *C. auris* strains had somewhat lower phosphomannan content than wild-type *C. albicans* SC5314 and slightly higher phosphomannan content than *mnn4*Δ/*mnn4*Δ *C. albicans*, but these differences were mostly not statistically significant (Table S4). Importantly, the positive (SC5314) and negative (*mnn4*Δ/*mnn4*Δ) controls show a highly significant difference (*p* = 3.89×10^-4^, Dunn’s test with Bonferroni correction) (Table S4).

Intriguingly, in micrographs from high-pressure freeze substitution transmission electron microscopy the parental B11220 (*FKS1-S639*) strain (Fig. 4A) has a significantly thicker inner cell wall and longer mannan fibrils (Fig. 4B; Tables S5, S6). However, none of the derived strains, including the CSP^S^ *FKS1-S639* isolate UACa140, differed from each other indicating that passaging on RPMI rather than amino acid status at residue 639 of Fks1 influenced cell wall ultrastructure (Fig. 4B; Tables S5, S6).

**Figure 4.**
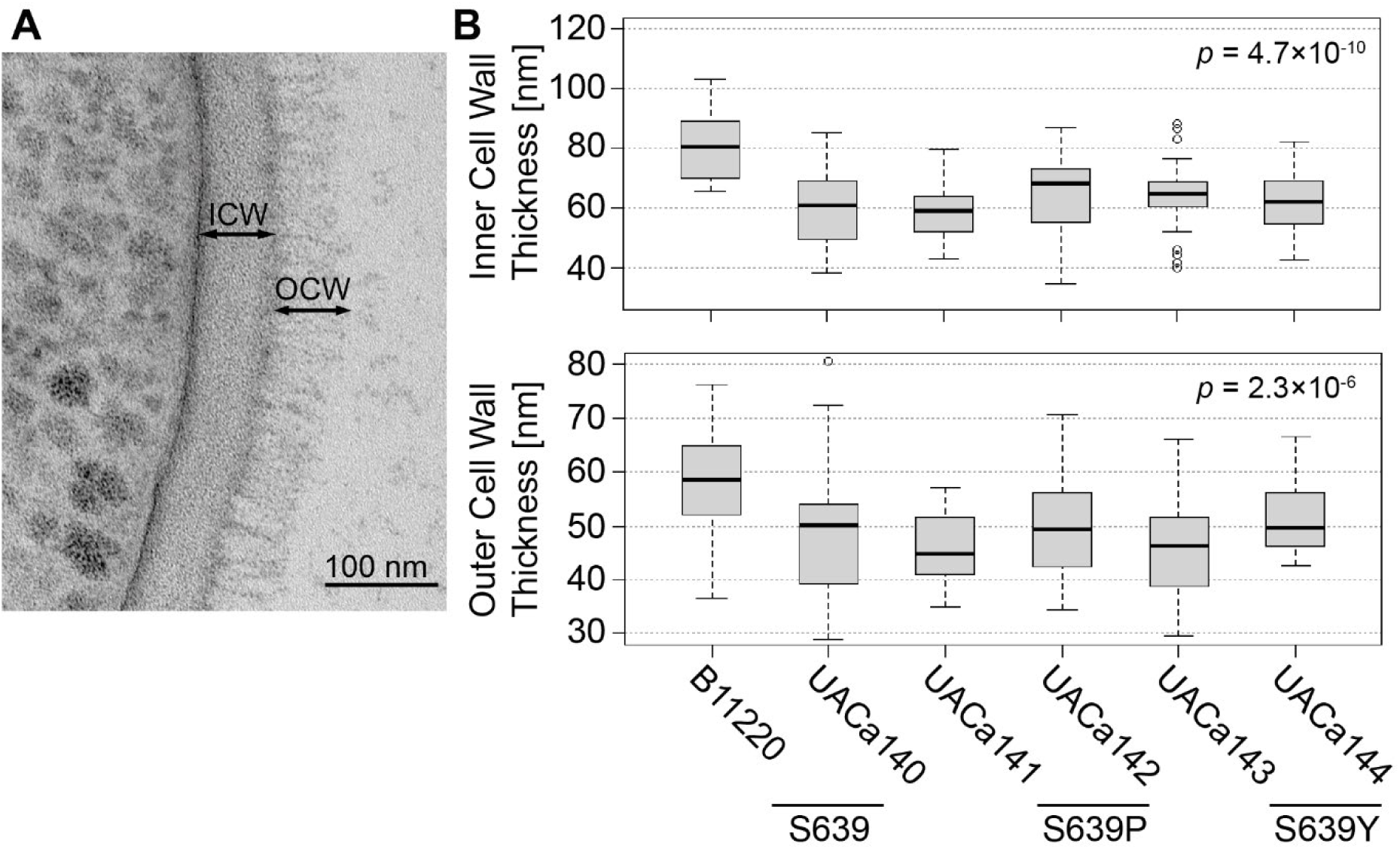
*In vitro* evolution of CSP resistance via point mutations in *FKS1* does not correlate with changes to the cell wall ultrastructure. Cells grown in RPMI were fixed by high-pressure freezing for transmission electron microscopy (TEM). (**A**) Representative cell wall TEM image of strain B11220; outer cell wall (OCW) and inner cell wall (ICW) are indicated. Scale bar represents 100 nm. (**B**) Top: box plot showing inner cell wall thickness [nm] of 30 cells from each of the indicated strains. Bottom: box plot showing length of mannan fibrils [nm] (outer cell wall thickness) of 30 cells from each of the indicated strains. *p*-values from a Kruskal-Wallis test, for *post-hoc* analysis with a Dunn’s test using a Bonferroni correction see Tables S5 & S6.

Taken together, this indicates that *FKS1* mutation in a *C. auris* clade II strain leads to a moderate increase in cell wall chitin and possibly a slight increase in cell wall phosphomannans but these changes barely affect cell wall ultrastructure during evolution of CSP resistance (Figs. 3, 4, S3).

### 3.5. Biofilm formation is not affected by *FKS1-S639* variant status

The dynamic regulation of cell wall glucan in *C. albicans* via exoglucanases, such as Xog1, determines the ability of cells to stick to each other and to surfaces which could influence biofilm formation (Tsai et al., 2011). Indeed, β-glucans contribute to biofilm biomass and to antimicrobial resistance of biofilms in *C. albicans* (Nett et al., 2007; Taff et al., 2012). Therefore, we tested whether the *FKS1-S639P/Y* variants have a different propensity to form biofilms. Neither biofilm biomass nor cell viability in biofilms was different in CSP^S^ *FKS1-S639* strains (B11220, UACa140) compared to the related CSP^R^ *FKS1-S639P/Y* strains (UACa141-UACa144) (Fig. 5). This shows that capability for biofilm formation is not altered in *FKS1- S639P/Y* variants of *C. auris* clade II strains.

**Figure 5.**
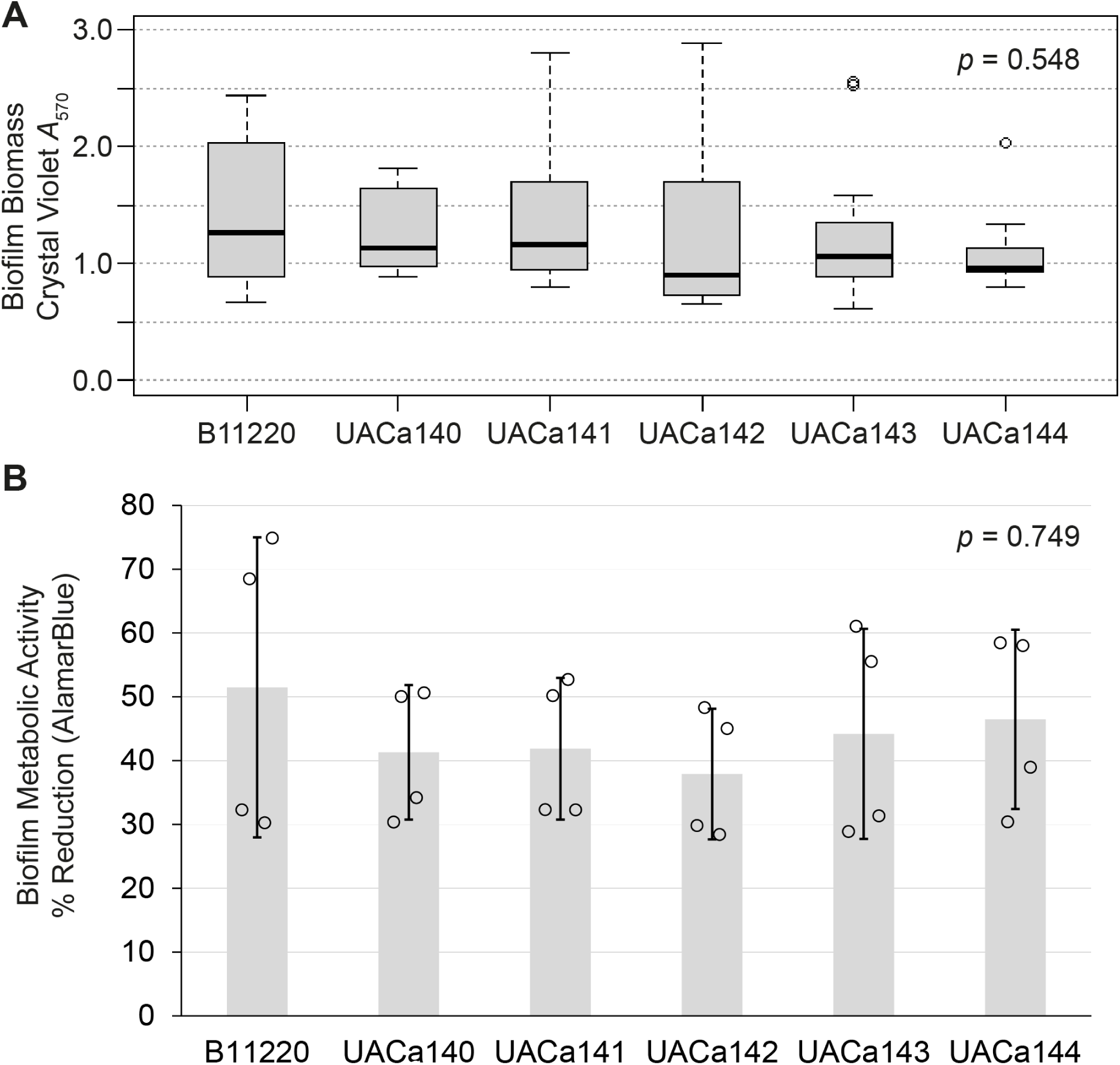
Biofilm mass and cell viability in biofilms is unaffected in *FKS1* variant strains. *p*-values were generated using a Kruskal-Wallis test. (**A**) Crystal Violet staining of biofilms produced by the indicated strains (n = 4 biological repeats each containing 3 technical repeats). Absorbance at 570 nm is shown in optical density units. (**B**) AlamarBlue staining of biofilms to measure metabolic activity of cells in biofilms is expressed as % reduction of resazurin into resofurin (Brown et al., 2022), bar chart shows average ± standard deviation as error bar (n = 4 biological repeats each containing 3 technical repeats).

### 3.6. *FKS1-S639* variant status does not impact virulence

Evolving resistance to environmental stresses, including antimicrobial treatment, produces a fitness trade-off which can result in attenuated virulence in both bacteria and fungi (Beceiro et al., 2013; Carolus et al., 2024b; Das et al., 2024; Geisinger and Isberg, 2017; Hill et al., 2015; Vincent et al., 2013). To test whether evolving resistance to CSP causes a trade-off with virulence, we employed the invertebrate infection model *Galleria mellonella* (Borman, 2022; Martinez et al., 2024). There were no statistically significant differences in killing *Galleria* larvae between CSP^S^ (*FKS1-S639*) and CSP^R^ (*FKS1-S639P/Y*) strains (Figs. 6, S5). This indicates that there is no trade-off between CSP-resistance mediated by an *FKS1* hotspot I variant and virulence in an invertebrate host for *C. auris* clade II strains. This stands in contrast to the observation of *C. albicans FKS1* variant strains being hypovirulent in *Drosophila* and mouse infection models (Ben-Ami et al., 2011). Although we do observe a fitness trade-off of CSP resistance evolution in *C. auris* leading to an increased susceptibility to some antifungal azole drugs and to cell wall stress (Fig. 2), this and subtle changes in cell wall composition do not seem impact the virulence of CSP^R^ strains at least in an invertebrate model of infection (Figs. 6, S5).

**Figure 6.**
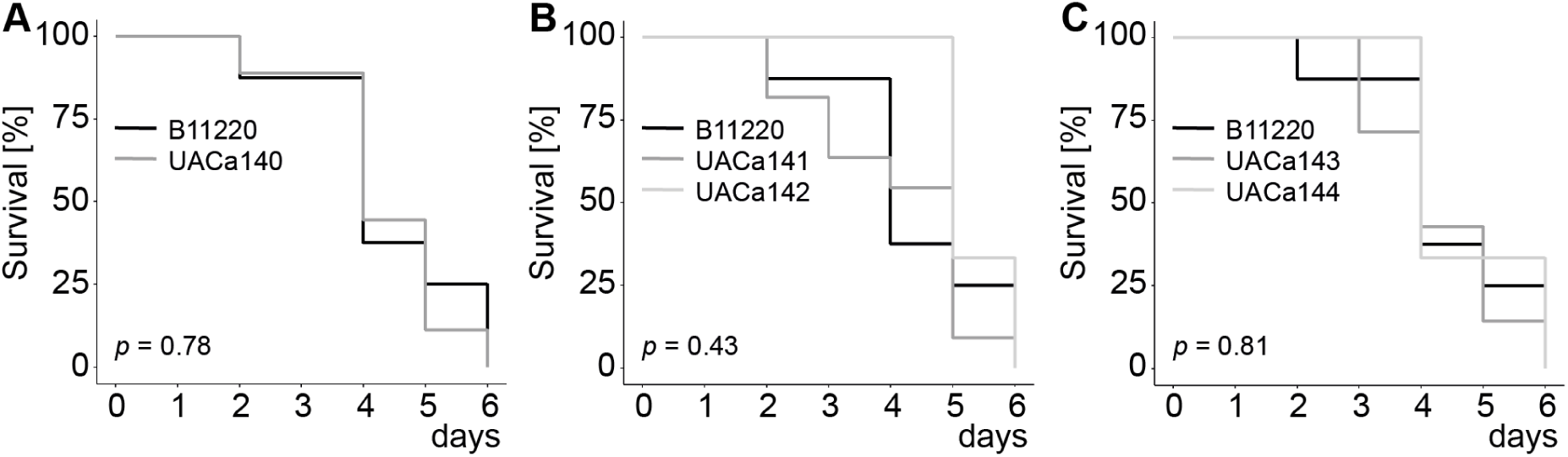
Virulence in the *G. mellonella* infection model is unaffected by *FKS1* variants. *C. auris* cells were grown in RPMI broth before inoculation into *G. mellonella* larvae. 20 larvae in 2 independent experiments were used to assess virulence, *p*-values were generated using a log-rank test. For a 6-ways comparison between all strains see Fig. S5. (**A**) Comparing parental B11220 and derived UACa140 strains, both carry serine at residue 639 of Fks1 and are CSP^S^. (**B**) Comparing parental B11220 (*FKS1-S639*, CSP^S^) and *in vitro* evolved CSP^R^ strains UACa141 and UACa142 strains (*FKS1-S639P*). (**C**) Comparing parental B11220 (*FKS1-S639*, CSP^S^) and *in vitro* evolved CSP^R^ strains UACa143 and UACa144 strains (*FKS1-S639Y*).

## 4. Conclusion

In *C. auris,* hotspot I mutations of *FKS1* leading to substitution of serine at residue 639 with phenylalanine, tyrosine, or proline (S639F,-Y,-P) in the Fks1 protein are known from clades I, III & IV isolates and have been linked to echinocandin resistance (Biagi et al., 2019; Chow et al., 2020; Chowdhary et al., 2018; Kordalewska et al., 2018). However, *C. auris* clade II strains had not previously been shown to evolve S639 *FKS1* hotspot I mutations in response to caspofungin exposure (Carolus et al., 2021; Hirayama et al., 2023). Intriguingly, *C. auris* clade II has considerable differences to the other clades in terms of genome structure (Muñoz et al., 2021), and its representatives rarely cause bloodstream infections and are often susceptible to antifungal drugs apart from fluconazole (Chow et al., 2020; Kwon et al., 2019; Welsh et al., 2019). These strands of evidence invite the speculation that clade II strains might be defective in developing echinocandin resistance.

Here, we demonstrate for the first time that a CSP^S^ strain (B11220) of the *C. auris* clade II can be *in vitro* evolved to CSP resistance by acquisition of *FKS1* hotspot I mutations using whole-genome sequencing (Fig. 1). These mutations lead to amino acid changes at residue 639 from serine to proline or tyrosine (Table 1). We subsequently examined these strains for cross-resistance & collateral sensitivity to antifungal drugs and cell wall stress. Changes in cell wall structure and biochemistry, biofilm formation capability, and virulence were also studied.

As anticipated, the *FKS1-S639P* and *FKS1-S639Y* variants conferred resistance not only to caspofungin but also to the echinocandins anidulafungin and micafungin (Fig. 2A). Notably, CSP^R^ strains exhibited collateral sensitivity to certain azole drugs (ICZ and PSZ) and increased susceptibility to cell wall stressors CFW and CR (Fig. 2). While there was a moderate increase in chitin content in CSP^R^ isolates, overall cell wall structure and phosphomannan levels remained unchanged (Figs. 3, 4). Additionally, biofilm formation was comparable across strains, and virulence in invertebrate models did not differ between strains with *FKS1* mutations and those without (Figs. 5, 6).

In conclusion, although the evolution of caspofungin resistance through *FKS1* mutations introduces some fitness trade-offs, it does not substantially compromise the pathogenic potential of this clade II *C. auris* isolate. This highlights the intricate relationship between antifungal resistance and virulence in this recently emerged fungal pathogen.

## Acknowledgements

We are grateful for technical support from the Centre for Genome-Enabled Biology and Medicine (CGEBM) of the University of Aberdeen, UK. *C. auris* strain B11220 was kindly provided by the Mycotic Diseases Branch, CDC (Atlanta, GA, USA), and *C. albicans* strains were kindly donated by Neil Gow. We thank Delma Childers for discussions.

## Transparency Declaration

The authors declare that there is no conflict of interest.

