## Supplementary Figures & Tables for "*In vitro* evolution of caspofungin resistance in *Candidozyma auris* via *FKS1* hotspot I mutations results in moderate fitness trade-offs but no reduction in virulence"

**Supplementary Table S1.** Strain list.

| Strain Name | Collection No. | Species | Relevant Genotype | Additional Information | Source |
|---|---|---|---|---|---|
| B11220 | UACa18 | *Candidozyma auris* | *MATα FKS1-S639* | clade II clinical isolate from auditory canal | (Lockhart et al., 2017) |
| | UACa140 | *Candidozyma auris* | *MATα FKS1-S639* | $CSP^S$ derivative of B11220 | this study |
| | UACa141 | *Candidozyma auris* | *MATα FKS1-S639P* | $CSP^R$ derivative of B11220 | this study |
| | UACa142 | *Candidozyma auris* | *MATα FKS1-S639P* | $CSP^R$ derivative of B11220 | this study |
| | UACa143 | *Candidozyma auris* | *MATα FKS1-S639Y* | $CSP^R$ derivative of B11220 | this study |
| SC5314 | UACa38 | *Candida albicans* | standard wild-type laboratory strain | | (Fonzi and Irwin, 1993) |
| Gow236 | UACa96 | *Candida albicans* | *mnn4 Δ /mnn4Δ::his-G-URA3-hisG* | | (Hobson et al., 2004) |

**Supplementary Table S2.** *p*-values from Tukey's HSD to assess differences between means of Calcofluor White mean fluorescence intensity measured by flow cytometry to estimate the chitin content of the cell wall between *Candidozyma auris* B11220, UACa140, UACa141, UACa142, and UACa143, and UACa144 in Figure 3A.

| comparison of strains | adjusted *p*-value |
|---|---|
| B11220 - UACa140 | 0.6138862 |
| B11220 - UACa141 | 0.1124297 |
| B11220 - UACa142 | 0.0065854 |
| B11220 - UACa143 | 0.0010180 |
| B11220 - UACa144 | 0.0001662 |
| UACa140 - UACa141 | 0.8488210 |
| UACa140 - UACa142 | 0.1563693 |
| UACa140 - UACa143 | 0.0288907 |
| UACa140 - UACa144 | 0.0045972 |
| UACa141 - UACa142 | 0.7229010 |
| UACa141 - UACa143 | 0.2482739 |
| UACa141 - UACa144 | 0.0510253 |
| UACa142 - UACa143 | 0.9458557 |
| UACa142 - UACa144 | 0.5124777 |
| UACa143 - UACa144 | 0.9469485 |

**Supplementary Table S3.** *p*-values from Tukey's HSD to assess differences between means of Calcofluor White mean fluorescence intensity measured by flow cytometry to estimate the chitin content of the cell wall between *Candidozyma auris* B11220, UACa140, UACa141, UACa142, UACa143, and UACa144 in Figure S3.

| comparison of strains | adjusted *p*-value |
|---|---|
| B11220 - UACa140 | 0.8888499 |
| B11220 - UACa141 | 0.0118693 |
| B11220 - UACa142 | 0.1389186 |
| B11220 - UACa143 | 0.0249510 |
| B11220 - UACa144 | 0.0171229 |
| UACa140 - UACa141 | 0.0702797 |
| UACa140 - UACa142 | 0.5702744 |
| UACa140 - UACa143 | 0.1430728 |
| UACa140 - UACa144 | 0.1003496 |
| UACa141 - UACa142 | 0.6825688 |
| UACa141 - UACa143 | 0.9973480 |
| UACa141 - UACa144 | 0.9999125 |
| UACa142 - UACa143 | 0.8957850 |
| UACa142 - UACa144 | 0.7993287 |
| UACa143 - UACa144 | 0.9999019 |

**Supplementary Table S4.** *p*-values from a Dunn's test with Bonferroni correction for multiple comparisons of means of the phosphomannan content of the cell wall [pg/cell] (Alcian Blue staining) between *Candidozyma auris* B11220, UACa140, UACa141, UACa142, UACa143 & UACa144, and *Candida albicans* SC5314 & *mnn4*Δ/*mnn4*Δ in Figure 3B.

| **comparison of strains** | **z-statistic** | **adjusted *p*-value** |
|---|---|---|
| B11220 - UACa140 | 0.440 | 1 |
| B11220 - UACa141 | 0.890 | 1 |
| B11220 - UACa142 | 0.795 | 1 |
| B11220 - UACa143 | 1.27 | 1 |
| B11220 - UACa144 | 1.57 | 1 |
| B11220 - SC5314 | 3.21 | 0.0376 |
| B11220 - *mnn4*Δ/*mnn4*Δ | -1.14 | 1 |
| UACa140 - UACa141 | 0.451 | 1 |
| UACa140 - UACa142 | 0.355 | 1 |
| UACa140 - UACa143 | 0.828 | 1 |
| UACa140 - UACa144 | 1.13 | 1 |
| UACa140 - SC5314 | 2.77 | 0.158 |
| UACa140 - *mnn4*Δ/*mnn4*Δ | -1.58 | 1 |
| UACa141 - UACa142 | -0.0958 | 1 |
| UACa141 - UACa143 | 0.378 | 1 |
| UACa141 - UACa144 | 0.682 | 1 |
| UACa141 - SC5314 | 2.32 | 0.575 |
| UACa141 - *mnn4*Δ/*mnn4*Δ | -2.03 | 1 |
| UACa142 - UACa143 | 0.473 | 1 |
| UACa142 - UACa144 | 0.778 | 1 |
| UACa142 - SC5314 | 2.41 | 0.444 |
| UACa142 - *mnn4*Δ/*mnn4*Δ | -1.93 | 1 |
| UACa143 - UACa144 | 0.304 | 1 |
| UACa143 - SC5314 | 1.94 | 1 |
| UACa143 - *mnn4*Δ/*mnn4*Δ | -2.41 | 0.451 |
| UACa144 - SC5314 | 1.63 | 1 |
| UACa144 - *mnn4*Δ/*mnn4*Δ | -2.71 | 0.188 |
| SC5314 - *mnn4*Δ/*mnn4*Δ | -4.35 | $3.89 \times 10^{4}$ |

**Supplementary Table S5.** *p*-values from a Dunn's test with Bonferroni correction for multiple comparisons of means of the inner cell wall (ICW) thickness measurements between B11220, UACa140, UACa141, UACa142, UACa143, and UACa144 in Figure 4B.

| comparison of strains | z-statistics | adjusted *p*-value |
|---|---|---|
| B11220 - UACa140 | -5.75 | $1.32\times10^{7}$ |
| B11220 - UACa141 | -6.40 | $2.31\times10^{9}$ |
| B11220 - UACa142 | -4.00 | $9.49\times10^{4}$ |
| B11220 - UACa143 | -4.62 | $5.77\times10^{5}$ |
| B11220 - UACa144 | -5.27 | $2.02\times10^{6}$ |
| UACa140 - UACa141 | -0.649 | 1 |
| UACa140 - UACa142 | 1.75 | 1 |
| UACa140 - UACa143 | 1.13 | 1 |
| UACa140 - UACa144 | 0.479 | 1 |
| UACa141 - UACa142 | 2.40 | 0.245 |
| UACa141 - UACa143 | 1.78 | 1 |
| UACa141 - UACa144 | 1.13 | 1 |
| UACa142 - UACa143 | -0.619 | 1 |
| UACa142 - UACa144 | -1.27 | 1 |
| UACa143 - UACa144 | -0.653 | 1 |

**Supplementary Table S6.** *p*-values from a Dunn's test with Bonferroni correction for multiple comparisons of means of the mannan fibril (outer cell wall, OCW) length measurements between B11220, UACa140, UACa141, UACa142, UACa143, and UACa144 in Figure 4B.

| comparison of strains | z-statistic | adjusted *p*-value |
|---|---|---|
| B11220 - UACa140 | -3.70 | $3.21\times10^{3}$ |
| B11220 - UACa141 | -5.21 | $2.84\times10^{6}$ |
| B11220 - UACa142 | -3.61 | $4.51\times10^{3}$ |
| B11220 - UACa143 | -4.57 | $7.31\times10^{5}$ |
| B11220 - UACa144 | -2.51 | 0.180 |
| UACa140 - UACa141 | -1.51 | 1 |
| UACa140 - UACa142 | 0.0867 | 1 |
| UACa140 - UACa143 | -0.868 | 1 |
| UACa140 - UACa144 | 1.19 | 1 |
| UACa141 - UACa142 | 1.59 | 1 |
| UACa141 - UACa143 | 0.639 | 1 |
| UACa141 - UACa144 | 2.70 | 0.105 |
| UACa142 - UACa143 | -0.955 | 1 |
| UACa142 - UACa144 | 1.10 | 1 |
| UACa143 - UACa144 | 2.06 | 0.594 |

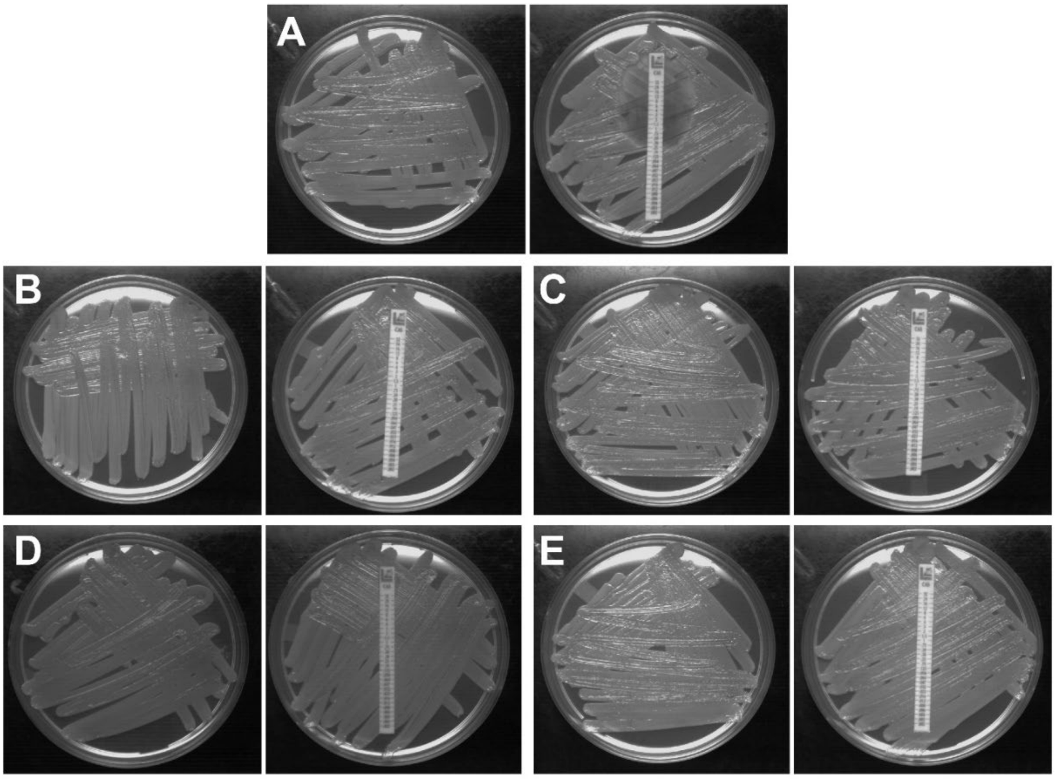


**Supplementary Figure S1. Growth controls (left) and caspofungin Etests (right) of B11220 passaged without caspofungin (A) or with caspofungin (B - E).** Isolates UACa140 (**A**), UACa141 (**B**), UACa142 (**C**), UACa143 (**D**), and UACa144 (**E**).

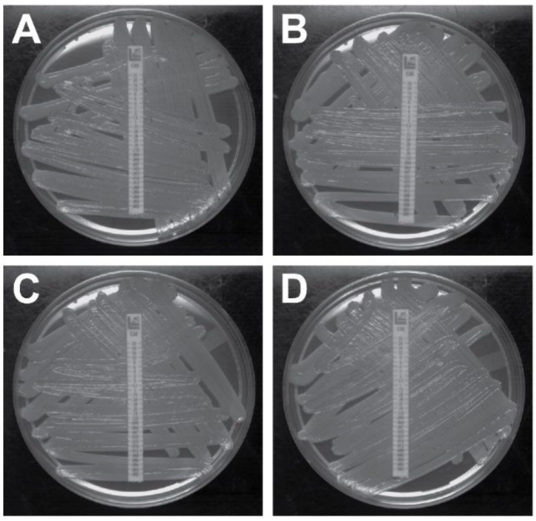


**Supplementary Figure S1. Caspofungin Etests performed on *in vitro* evolved caspofungin - resistant isolates after growth without caspofungin selection pressure for 2 weeks.** Isolates UACa141 (**A**), UACa142 (**B**), UACa143 (**C**), and UACa144 (**D**).

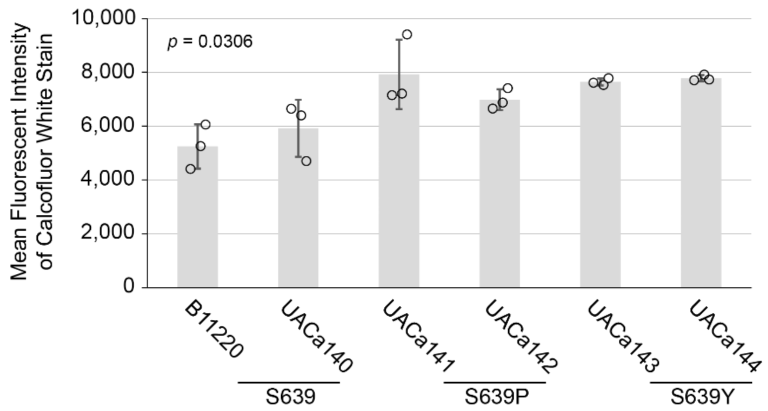


**Supplementary Figure S3. Cells of CSP$^R$ strains have a higher chitin content.** (**A**) Bar chart showing mean fluorescent intensity of Calcofluor White staining of the indicated *C. auris* strains determined by flow cytometry (n = 3, ~50,000 cells measured in each sample) *p*-value from an ANOVA, for *post-hoc* analysis with Tukey's HSD see Table S3. An independent run is shown in Fig. 3A.

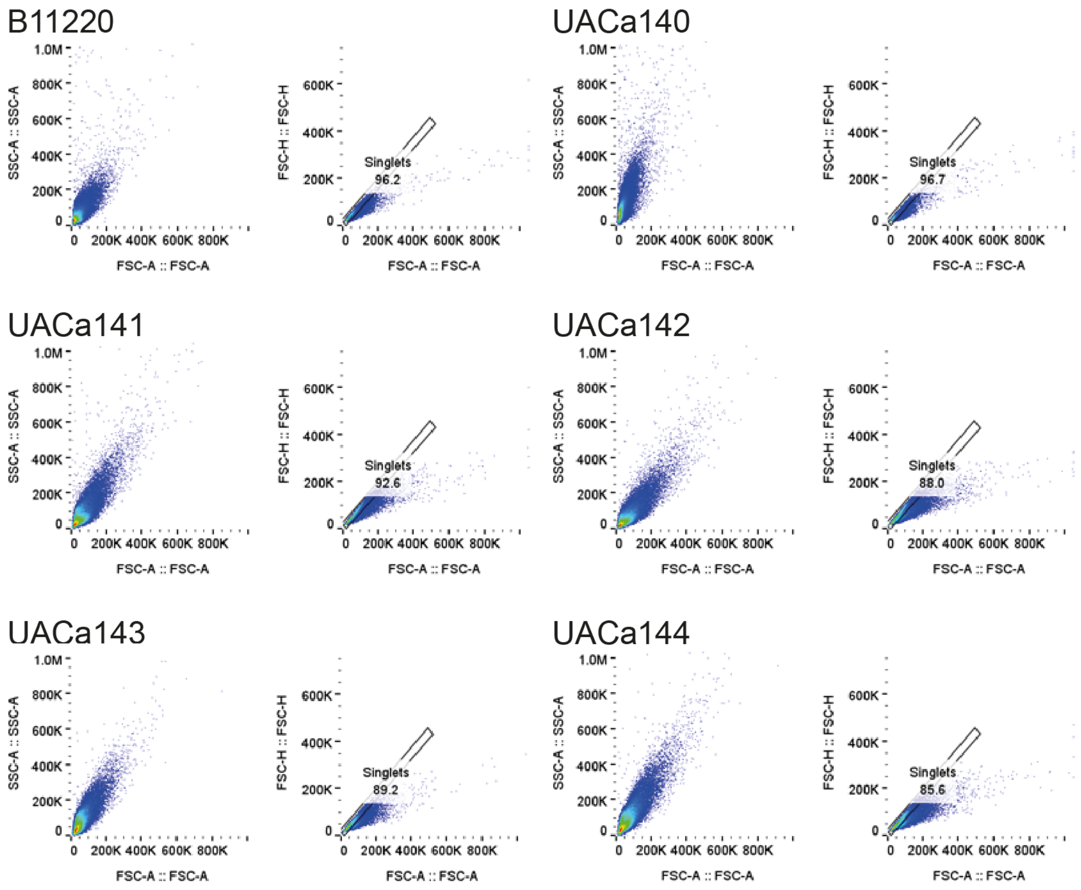


**Supplementary Figure S4. Representative scatter plots showing gating strategy during flow cytometry.** A representative example of forward and side scatter to illustrate the gating strategy from the experiment in Fig. 3A is shown for each strain.

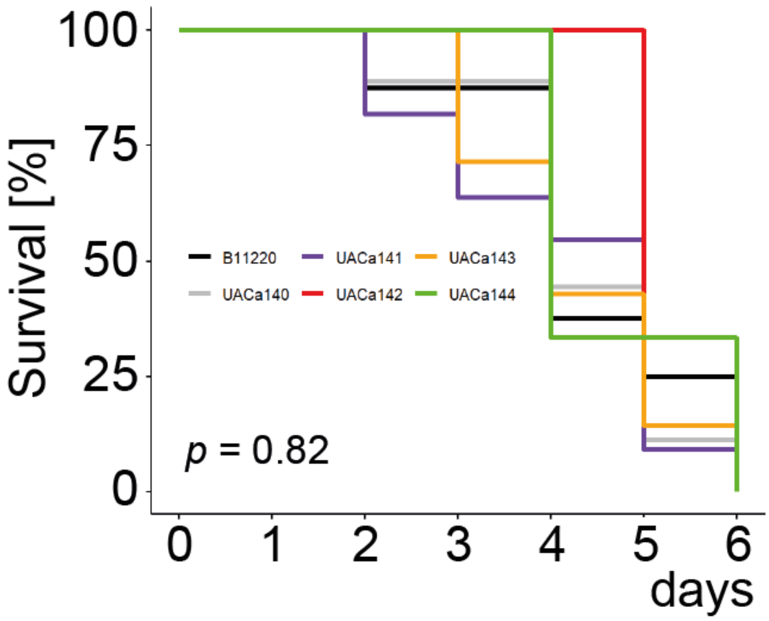


**Supplementary Figure S5. Virulence in the *G. mellonella* infection model is unaffected by *FKS1* variants.** *C. auris* cells were grown in RPMI broth before inoculation into *G. mellonella* larvae 20 larvae in 2 independent experiments were used to assess virulence, *p* - value was generated using a log - rank test. Comparing parental $CSP^S$ B11220 and derived $CSP^S$ UACa140 strains (both carrying a serine at residue 639 of Fks1) with *in vitro* evolved $CSP^R$ strains UACa141 & UACa142 (*FKS1 - S639P*) and UACa143 & UACa144 (*FKS1 - S639Y*).
